# Next-generation snow leopard population assessment tool: multiplex-PCR SNP panel for individual identification from feces

**DOI:** 10.1101/2024.09.19.613565

**Authors:** Katherine A. Solari, Shakeel Ahmad, Ellie E. Armstrong, Michael G. Campana, Hussain Ali, Shoaib Hameed, Jami Ullah, Barkat Ullah Khan, Muhammad A. Nawaz, Dmitri A. Petrov

## Abstract

Snow leopards, *Panthera uncia*, are under threat from numerous pressures and are the focus of a great deal of conservation efforts. However, their elusive nature makes it difficult to estimate population sizes. Current methods used to monitor local population sizes include visually identifying individuals from camera trap photos and genetically identifying individuals from fecal samples using microsatellite loci. Here, we present a new method for identifying snow leopard individuals from fecal samples using a multiplex PCR single nucleotide polymorphism (SNP) panel method. The SNP panel we present consists of 144 SNPs and utilizes next-generation sequencing technology, making it cheaper and easier than current microsatellite methods. We validate our SNP panel with paired tissue and fecal samples from zoo individuals, showing a minimum of 96.7% accuracy in allele calls per run. We then generate SNP data from 235 field-collected fecal samples from across Pakistan to show that the panel can reliably identify individuals from low-quality fecal samples of unknown age and is robust to contamination. We also show that our SNP panel has the capability to identify first-order relatives and provides insights into the geographic origin of samples. This SNP panel will empower the snow leopard research community in their efforts to assess local and global snow leopard population sizes. More broadly, we present a method for developing a SNP panel that utilizes open source software for SNP selection and primer design, Illumina sequencing technology, and a streamlined lab and bioinformatics protocol which can be used to create similar SNP panels for any species of interest for which adequate genomic reference data is available.

## Introduction

Snow leopards (*Panthera uncia*), are an enigmatic cat occupying mountainous areas of 12 South and Central Asian countries. They are notoriously difficult to study due to their harsh habitat, extremely elusive behavior, and low population densities (Bian et al., 2023; Chetri et al., 2019; R. K. Sharma et al., 2021). As such, lack of knowledge is often cited as one of the main issues hindering snow leopard conservation (McCarthy & Mallon, 2016). Although currently listed as Vulnerable by the IUCN (Ale & Mishra, 2018), snow leopards were previously listed as Endangered and are in need of protection as they are under threat from numerous pressures including habitat loss (Khan et al., 2021), declines in natural prey (Lovari et al., 2009), retaliatory killings due to livestock predation (Jackson et al., 2010), climate change (J. Li et al., 2021), and poaching (Aryal, 2017; Network, 2014).

One of the key knowledge gaps currently limiting snow leopard conservation efforts is an understanding of local and global population size. A dedicated international initiative was established to fill this important gap in 2017 – Population Assessment of the World’s Snow Leopards (PAWS) (K. Sharma et al., 2024). Up to this point, researchers have utilized extensive camera trapping efforts (e.g., (Jackson et al., 2006)), where individuals are identified based on unique coat patterns, as well as fecal microsatellite methods to identify individuals and estimate local population sizes (Chetri et al., 2019; Janečka et al., 2008, 2011; Karmacharya et al., 2011). However, in the case of camera trapping, identification errors are underappreciated and make it difficult for different datasets to be directly comparable (Johansson et al., 2020; Wong & Kachel, 2024). In the case of microsatellites, it can be difficult to standardize methods among labs which hinders the ability to directly compare datasets processed by different groups (Ellis et al., 2011). Microsatellite methods can also be very expensive to run on fecal samples since multiple replicates are necessary (Bayes et al., 2000) and extremely time-consuming as datasets are usually called manually.

Given the limitation of microsatellite methods and the exponentially decreasing cost of next-generation sequencing (NGS), the snow leopard community stands to benefit from a SNP panel for individual identification that can utilize NGS technology (Janjua et al., 2020). Specifically, the ideal SNP panel for individual identification would capitalize on inexpensive and flexible short-read sequencing (such as Illumina), work on low-quality fecal samples of unknown age, and employ a straightforward lab and bioinformatics protocol that can be run in snow leopard range countries without the need for overly specialized equipment.

The use of SNP panels for genotyping animals through non-invasive fecal samples has expanded over the last few years, with many different methods being utilized. For example, single primer enrichment technology through Allegro Targeted Genotyping has been used in feral horses (Gavriliuc et al., 2022), Kompetitive allele specific PCR (KASP) assays in forest elephants (Bourgeois et al., 2019), Fluidigm assays in mountain lions (Buchalski et al., 2022) and bison (Wehrenberg et al., 2024), MassARRAY systems in bats (Thavornkanlapachai et al., 2024), DNA hybridization capture in coyotes and foxes (Parker et al., 2022), and multiplex PCR (mPCR) methods including Genotyping-in-Thousands by sequencing (GT-seq) in deer (Burgess et al., 2022), polar bears (Hayward et al., 2022) and tigers (Natesh et al., 2019). However, many of these methods rely on specialized and expensive reagents or pieces of equipment, do not utilize Illumina sequencing technology for genotyping, and/or necessitate the use of service providers for primer design. Both Fluidigm and KASP assays detect SNPs by utilizing fluorescently labeled allele-specific primers with a fluorescence-based genotyping system (He et al., 2014). MassARRAY utilizes locus-specific PCR reaction followed by a single mass-modified base extension at the site of interest and a mass spectrometry-based genotyping system. DNA hybridization capture enriches areas of interest using probes and utilizes Illumina sequencing; however, this procedure employs very expensive reagents and a long, complex lab protocol. Allegro Targeted Genotyping and GT-seq both utilize Illumina sequencing for reporting; however, in both cases, external companies are required for primer design services.

Here, we develop a snow leopard SNP panel that consists of 144 SNPs and utilizes widely available Illumina sequencing technology, open-source software for SNP selection and primer design, and a streamlined lab and bioinformatics protocol that does not require specialized equipment. We validate our SNP panel with paired tissue and fecal samples from zoo individuals and subsequently apply the panel to 235 field-collected fecal samples from across the snow leopard range in Pakistan to show that it can reliably identify individuals from low quality fecal samples.

In doing this, we have achieved three main objectives: 1) we have presented an affordable protocol that can be used to create a SNP panel for any species of interest for which adequate genomic reference data is available; 2) we have created and validated a SNP panel for snow leopard individual ID from fecal samples that can be immediately used by the snow leopard research community in their efforts to assess local and global snow leopard population sizes; and 3) we have explored what other insights this snow leopard SNP panel can offer and have shown that it also has the capability to identify first-order relatives as well as general geographic origin.

## Methods

### Sample collection

A total of 15 fecal samples were collected from five snow leopards at the San Francisco Zoo (SF Zoo). These samples were collected within a day of being deposited and were stored in ziplock bags in a -80°C freezer until processing. The five SF Zoo snow leopards sampled and their relationship to each other are indicated in Table 1. Fecal samples were also collected from across the snow leopard range of Pakistan between 2018 and 2023. A line transect method was used for fecal sample collection and samples were collected into 50mL tubes with 30mL of silica beads. Information including location, habitat type, and estimated age of scat was recorded. A total of 1,174 putative snow leopard fecal samples were collected from across the snow leopard range in Pakistan covering three major mountain ranges – Himalaya, Hindu Kush, and Karakoram, each falling in a different administrative regions of Pakistan – District Chitral of Khyber Pakhtunkhwa province, Gilgit-Baltistan, and Neelum Valley of Azad Jammu and Kashmir, respectively. Of these 1,174 samples, only 42 (3.6%) were considered to be fresh, 1,091 samples (93%) were estimated to be over a week old and the majority of these, 807 samples (69%), were estimated to be at least two weeks old.

**Table 1.**
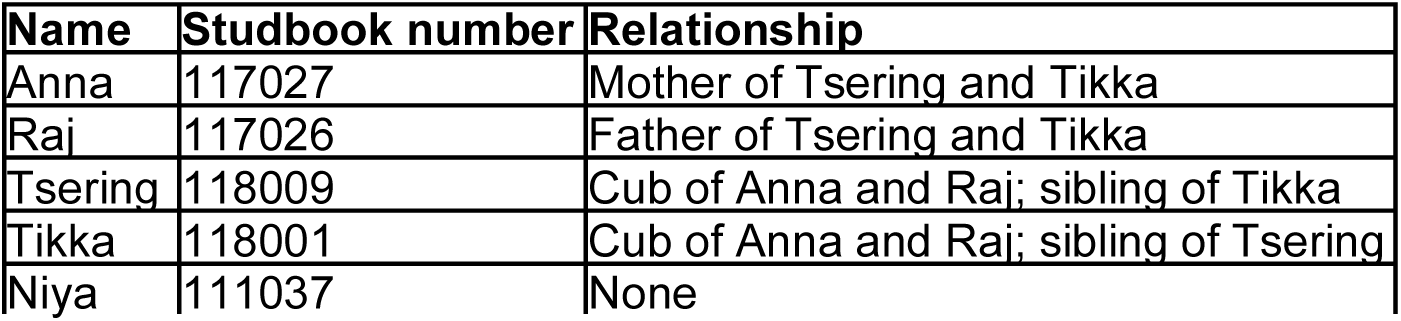
Relatedness among San Francisco Zoo samples.

### DNA extraction

We extracted DNA from all fecal samples using the Zymo Quick-DNA Fecal/Soil Microbe Miniprep Kit (catalog # D6010) following the manufacturer’s guidelines with minor adjustments. In order to prioritize the recovery of host DNA, we scraped the outside of each fecal sample with a sterile razor blade and added <150 mg (pea-size amount) of this material to 2mL beadbashing tubes for DNA extraction. SF Zoo fecal samples were processed frozen and Pakistan fecal samples, which were stored dried in silica beads, were processed at room temperature. Due to the dryness of the Pakistan samples, we added 900uL of beadbashing buffer (rather than the 750μL indicated in the protocol) to each sample in order to ensure 400μL of liquid could be recovered. We homogenized fecal samples in a BioSpec mini beadbeater 96 (catalog # NC0141170) for five minutes. We conducted all fecal extractions in a hood used only for fecal sample extractions and we treated the fecal extraction hood with 10% bleach, 70% ethanol, and UV light between each set of extractions. We included an extraction null in every set of fecal extractions to monitor for contamination.

### Species identification

We conducted species identification on all of the Pakistan fecal samples using universal mammal primers (MiMmamal-U) developed by Ushio et al. 2017 (Ushio et al., 2017) which amplifies ∼171 bp of the mitochondrial 12S rRNA gene. We employed a two-step PCR protocol so that samples could be uniquely barcoded and sequenced in pools. Each PCR1 reaction consisted of 5μL of AmpliTaq Gold 360 Master Mix, 2μL of water, 0.5μL of BSA, and 0.5μL of each of the forward and reverse primer at 10mM concentration and 2μL of extracted DNA. Primers used in PCR1 were the MiMammal-U primers from Ushio et al. 2017 (Ushio et al., 2017) followed by a spacer of six Ns and a sequencing primer (Forward - TCGTCGGCAGCGTCAGATGTGTATAAGAGACAGNNNNNNGGGTTGGTAAATTTCGTGCCA GC; reverse - GTCTCGTGGGCTCGGAGATGTGTATAAGAGACAGNNNNNNCATAGTGGGGTATCTAATCCCAGTTTG). The PCR1 cycle consisted of 10 minutes at 95°C, followed by 32 cycles of 30 seconds at 95°C, 30 seconds at 60°C and 30 seconds at 72°C, and completed with 7 minutes at 72°C and a 4°C hold.

Each PCR2 reaction consisted of 5μL of AmpliTaq Gold 360 Master Mix, 2.5μL of water, 0.5μL of each of the forward and reverse primers at 10mM concentration and 1.5μL of PCR1 product. The PCR2 primers were Illumina Nextera unique dual indexing primers ordered from Integrated DNA Technologies (IDT) consisting of an Illumina flow cell adapter sequence, a 10bp unique barcode, and a sequencing primer. We added a unique set of barcodes to each sample. The PCR2 cycle consisted of 10 minutes at 95°C, followed by 12 cycles of 30 seconds at 95°C, 30 seconds at 60°C and 10 seconds at 72°C, and completed with 7 minutes at 72°C and a 4°C hold. We included a PCR1 and PCR2 null with every sample set.

We took numerous precautions to limit contamination. We prepared PCR1 reactions in a pre-PCR hood that was treated with 10% bleach and UV light between each use and then ran PCRs in a different laboratory space on a separate floor of the building. We also prepared PCR2 reactions in a pre-PCR hood; however, we added the template, in this case the PCR1 product, in the post-PCR laboratory. We maintained a unidirectional flow of materials such that nothing from the post-PCR lab space came back to the pre-PCR lab space. We bleached PCR machines between each use and sprayed tubes containing PCR1 product with 10% bleach on the outside before opening during the addition of PCR1 product to the PCR2 reaction. We ran subsets of PCR1 and PCR2 products on a 1% agarose gel to monitor contamination and we ran a negative control with each PCR1 and PCR2 reaction. We pooled 2μL of PCR2 product from each sample and cleaned this pool using a Zymo clean and concentrator-25 kit. We sequenced samples on a MiSeq machine using a 2x250 bp configuration.

We removed MiMammal primers using Cutadapt v.4.4 (Martin, 2011) and retained only reads with the correct primer sequences. We then paired and merged reads using PEAR v0.9.6 (Zhang et al., 2014) and concatenated all of our sequence data and removed chimeras using -*- uchime3_denovo* in VSEARCH v2.22.1 (Rognes et al., 2016). In VSEARCH, we merged identical sequences using *--derep_fulllength* and clustered remaining sequences with pairwise identity of 0.97 or higher using *--cluster_size*. This resulted in the final list of unique sequences present in our dataset, each representing a unique operational taxonomic unit (OTU). We then calculated the number of exact full-length matches to each OTU for each sample using VSEARCH --*search_exact*.

We blasted each OTU sequence present in our dataset to the nt database using ‘blastn’ from ncbi-blast+ v.2.11.0 (Camacho et al., 2009) to identify matching sequences with a percent identity cut off of 93% and added taxon information using EFetch v.16.2 (part of NCBI’s E-utilities). We considered all samples with over 50 reads after all filtering and with over 51% of those reads assigned to snow leopard to be snow leopard scat (Table S1).

### SNP panel design

We selected SNPs from a previously published genomic dataset consisting of 37 snow leopards (Solari et al., 2023). We used a preliminary version (v.0.1.0) of the mPCRselect pipeline (Armstrong, Li, et al., 2024) to select SNPs to include in the SNP panel. First, we only considered SNPs on scaffolds corresponding to known autosomes and in regions of the genome with a mappability score of one as designated by genmapv1.3.0 (Pockrandt et al., 2019) using flags “-K 30” and “-E 2”. We then removed SNPs with a mapping quality (MQ) of less than 20, singletons, SNPs that had any variable sites (SNPs or Indels) within 40 bp up or downstream, indels, and non-biallelic SNPs. We then thinned the remaining SNPs to only retain a set of SNPs that were separated by 100,000 bp or more to roughly filter for SNPs that were in linkage disequilibrium.

Solari et al. (2023) identified four genetically distinct groups in their WGS dataset – North, consisting of Mongolia and Russia; Captive/Kyrgyzstan, consisting of the zoo population and Kyrgyzstan; South, consisting of Pakistan, Afghanistan, and Tajikistan; and India, consisting of a single sample from India. We used this information on genetically distinct groups to identify the 250 SNPs with the highest pairwise *F*_ST_ between the four groups using VCFtools *--weir-fst-pop* (Danecek et al., 2011) as well as the 250 SNPs with the highest within population pi for each group with more than one representative (India was excluded due to small sample size) in the reference dataset (North, Captive/Kyrgyzstan, and South) using the VCFtools *--site-pi*. We will refer to these SNP categories as *F*_ST_, North, Captive/Kyrgyzstan, and South, respectively.

We input these 1,000 target SNPs into NGS-PrimerPlex (Kechin et al., 2020) for compatible primer design with the following parameters: “-ad1 TCGTCGGCAGCGTCAGATGTGTATAAGAGACAG -ad2 GTCTCGTGGGCTCGGAGATGTGTATAAGAGACAG -minampllen 26 -optampllen 50 - maxampllen 120 -maxprimerlen 26 -minprimerlen 18 -blast -skip -primernum1 25 -tries 500”. Out of the 1000 SNPs, we had to remove 81 SNPs because the program reported that primers could not be designed for these regions and then we ran the remaining 919 SNPs again.

NGS-PrimerPlex output 144 primer pairs that could be pooled together (37 *F*_ST_ SNPs, 45 North SNPs, 31 Captive/Kyrgyzstan SNPs, and 31 South SNPs). We ordered these 144 primer pairs, with sequencing primers attached, from IDT (Table S2). We ordered primers dry and reconstituted them to a 100μM stock solution using IDTE buffer (pH 8). We pooled an equal amount of each 100μM primer to make a stock pool which was diluted with water to make a 10μM working stock primer pool.

### SNP panel amplification

Similarly to species identification, we also employed a two-step PCR protocol for the SNP panel amplification procedure. Each PCR1 reaction consisted of 10μL of 2X MasterMix from Qiagen Multiplex PCR Plus Kit, 1μL of water, 1μL of BSA, 4μL of the 10uM primer pool and 4μL of extracted DNA. The PCR1 cycle consisted of 5 minutes at 95°C, followed by 40 cycles of 30 seconds at 95°C, 90 seconds at 60°C and 30 seconds at 72°C, and completed with 10 minutes at 68°C and a 4°C hold. Each PCR2 reaction consisted of 4μL of 2X MasterMix from Qiagen Multiplex PCR Plus Kit, 3μL of water, 1μL of each of the forward and reverse barcoding primers at 10mM concentration and 1μL of PCR1 product. The PCR2 primers were the same primers used in PCR2 of the species identification procedure – Illumina Nextera unique dual indexing primers ordered from IDT. The PCR2 cycle consisted of 5 minutes at 95°C, followed by 12 cycles of 30 seconds at 95°C, 90 seconds at 62°C, and 30 seconds at 72°C, and completed with 10 minutes at 68°C and a 4°C hold. We included a PCR1 and PCR2 null in each set of reactions. We ran the SNP panel on each confirmed snow leopard sample from Pakistan between two to four times (as indicated in Table S3), two times on each of the SF Zoo fecal samples, and at least twice on every extraction null.

As our species identification results display, snow leopard scat can be confused with scat from other species. If the SNP panel is mistakenly run on a scat sample from a different species, it is important that we are able to distinguish SNP panel results from non-snow leopards such that they are not mistakenly included in the dataset. To ensure that SNP panel results from different species are easily identifiable, we ran the SNP panel on four samples from each of the three other species most commonly represented in the Pakistan fecal sample set - fox (*Vulpes spp.*), dog/wolf (*Canis lupus*), and lynx (*Lynx lynx*), as well as the additional feline species found in the sample set - Asian leopard (*Panthera pardus*) and leopard cat (*Prionailurus bengalensis*). We used four samples confidently identified as coming from each of these other species based on species identification sequencing results (Table S1). We did not run these samples in replication.

In order to limit contamination, we took the same anti-contamination steps described above for the species identification two-step PCR protocol. However, in this case, we did not visualize any PCR1 or PCR2 products on a gel as this was an additional opportunity for contamination and we found gel visualizations to be uninformative regarding amplification success. We provide additional details regarding steps taken to limit contamination in the supplementary methods.

Similarly to the species identification protocol, we pooled 2μL of PCR2 product from each sample and cleaned it using a Zymo clean and concentrator-25 kit. We sequenced samples on the Illumina NovaSeq X Plus using 2X150 bp configuration. A detailed wet lab protocol for running the SNP panel is provided in the Supplementary (Supplementary File 1).

### Variant calling

We mapped Illumina reads to the snow leopard reference genome (Armstrong et al., 2022) using BWA-MEM v.0.7.17 (H. Li, 2013). We sorted and indexed resulting bam files using SAMtools v.1.16.1 (H. Li et al., 2009). We removed reads with a MQ of zero using SAMtools *view* with the flags “-h -q 1”. We generated genotype likelihoods in target amplicon regions using BCFtools *mpileup* with the flags “-A -a AD,DP” and the “-R” flag followed by a bed file of target amplicons. We then called variants using BCFtools *call* with the flags “-m -Oz -f GQ”. We then filtered the data to only include target SNPs that had a depth ≥6, and a genotype quality score ≥20 using VCFtools and the flag “--bed” followed by a bed file containing target SNP locations, the flag “--minDP 6”, and the flag “--minGQ 20”, respectively.

### SNP panel performance assessment

We calculated the total number of raw reads and the number of reads remaining after filtering for MQ using SAMtools *flagstat*. In order to assess how many reads were mapping to the target amplicon regions, we used BEDtools *intersect* to create a subset of the bam files (post MQ filtering) consisting of only the 144 target regions with a 50bp buffer on either side and calculated read numbers using SAMtools *flagstat*. We calculated the depth of coverage for each SNP in each run after removing reads that mapped with a MQ of zero using BEDtools *multicov* (Quinlan & Hall, 2010).

We calculated the number of SNPs successfully genotyped in each run using VCFtools *- -missing-indv*. This allowed us to assess variation in SNP panel success between different runs of the same sample.

We then created a separate VCF file for each sample group (Pakistan n=604, SF Zoo n=30, Other species n=20, Nulls n=121) using VCFtools with the flag “-keep” followed by a list of samples to be kept in each group. We calculated allele frequencies within each group of samples using VCFtools --*freq*. We calculated the heterozygosity of each run using BCFtools *stat* with the flag “-S” followed by a text file with a list of all the samples.

### Calculating genotyping accuracy and allelic drop out with paired WGS data

High coverage (>40X) WGS data was available for four of the five SF Zoo individuals for which we obtained fecal samples (Armstrong, Carey, et al., 2024). We mapped WGS data to the snow leopard reference genome (Armstrong et al., 2022) using BWA-MEM v.0.7.17 (H. Li, 2013). We sorted and indexed resulting bam files using SAMtools v.1.16.1 (H. Li et al., 2009). We calculated the mean depth of coverage for each sample using SAMtools (Table S4). We called SNPs in target amplicon regions using BCFtools *mpileup* with the flags “-A -a AD,DP” and the “-R” flag followed by a bed file of target amplicons. We followed this by BCFtools *call* with the flags “-m -Oz -f GQ”. We then filtered SNP calls to only include the target SNPs using VCFtools and the flag “--bed” followed by a bed file containing target SNP locations.

We manually calculated genotyping accuracy and allelic dropout rate for all of the fecal samples from the four zoo individuals for which we had WGS data to serve as ‘true’ genotypes. We only retained mPCR runs that yielded genotypes for 20 or more SNPs. Genotyping accuracy at the SNP level for each mPCR run was calculated as the percent of successfully genotyped SNPs that were the same genotype as the high-coverage WGS data. Genotyping accuracy at the allele level was calculated as the percent of genotyped alleles that matched alleles called from WGS data. Allelic dropout rate for each mPCR run was calculated by dividing the number of SNPs with allelic dropout (SNPs that were identified at homozygous in mPCR results but were in fact heterozygous based on high-coverage WGS data) by the total number of heterozygous SNP for that individual (calculated from WGS high-coverage data) that were successfully genotyped in that mPCR run (Table S5).

### Identifying individuals from mPCR output

We only kept mPCR runs that were successfully genotyped at 20 or more SNPs. We then calculated the number of shared alleles between all sample pairs in each sample set using Plink v.1.90b5.3 with the flags “--allow-extra-chr --double-id --genome full”. This conducts a pairwise comparison between each possible pair of runs and calculates the number of SNPs identical by state (IBS) at both alleles (IBS2), at one allele (IBS1), and at neither allele (IBS0). We used this data to calculate the proportion of alleles in IBS as P(IBS2) + 0.5*P(IBS1). We considered two samples to be the same individual if they had 95% or more alleles IBS across at least 20 SNPs (Table S6).

As a way of validating our individual ID results for the Pakistan samples, we visualized how close samples from the same individual were to each other in geographic space compared to samples from different individuals. Here, our assumption was that samples collected from the same individual would be more likely to be close to each other than samples from two different individuals. To do this, we used Geographic Distance Matrix Generator (Ersts, n.d.) to calculate the physical pairwise distance between each sample pair from gps coordinates. We then computed an Empirical Cumulative Distribution Function (ECDF) for sample pairs representing the same individual versus sample pairs from different individuals using the *ecdf* function in R v.4.1.2 (R Core Team, 2020).

### Assessing relatedness among samples

Out of the five SF Zoo individuals, four consisted of a family unit of two parents and two offspring. We identified these first-order relationships and compared the proportion of alleles IBS in these related pairs to the proportion of alleles IBS in non-first-order pairs.

To get a sense of whether or not the SNP panel was able to detect relatedness among the Pakistan samples, we assessed the correlation between the proportion of alleles IBS and geographic distance between samples. Here, our assumption is that first degree relatives are more likely to be in closer proximity to each other than to non-first degree relatives. Little is known about snow leopards’ dispersal behavior, but it is known that cubs likely stay with their mother for almost two years (Johansson et al., 2021). We used the same geographical pairwise distances between sample pairs used in the ECDF analysis, but only pairwise comparisons between samples from different individuals were considered. We compared the proportion of allele IBS and geographic distance between sample pairs using Pearson correlation coefficient calculated using the ggpubr v.0.4.0 (Kassambara, 2020) package in R.

### Geographic assignment

Geographic assignment was a secondary goal of the SNP panel. Most SNPs were targeting genetic variation within populations for individual ID (using pi within population); however, 37 SNPs in this panel targeted genetic variation between populations with the aim of informing general geographic assignment. In order to assess how informative our SNP panel was for this purpose, we conducted a principal component analysis (PCA) of genotype data from the fecal samples combined with wild samples from across the range. To do this, we combined the VCF file for all of the fecal samples with a VCF file of just the 144 target SNPs for the range-wide WGS data from Solari et al. 2023 using the ‘vcf-merge’ command from VCFtools. We then conducted a PCA using Plink v.1.90b5.3 and visualized the results in R using ggplot.

## Results

### Pipeline development

Our goal was to not only develop a SNP panel to be used on snow leopard fecal samples, but to establish a protocol using only open-source software that could be utilized to develop a SNP panel for any species (Fig. 1). We utilized standard methods for DNA extraction and species ID (Fig. 1, blue). We then applied new SNP selection criteria (Armstrong, Li, et al., 2024) and employed an open-source and under-utilized primer design software, NGS-PrimerPlex (Kechin et al., 2020), to create what we show to be a highly-effective SNP panel (Fig. 1, green). Lastly, we utilized a streamlined two-step PCR approach for sample processing, the most cost-effective Illumina sequencing technology for data generation, and standard open-source software for data processing (Fig. 1, teal). The protocol that we outline can be easily replicated to create a SNP panel for any species with the only limiting factors being available genomic reference data.

**Fig. 1.**
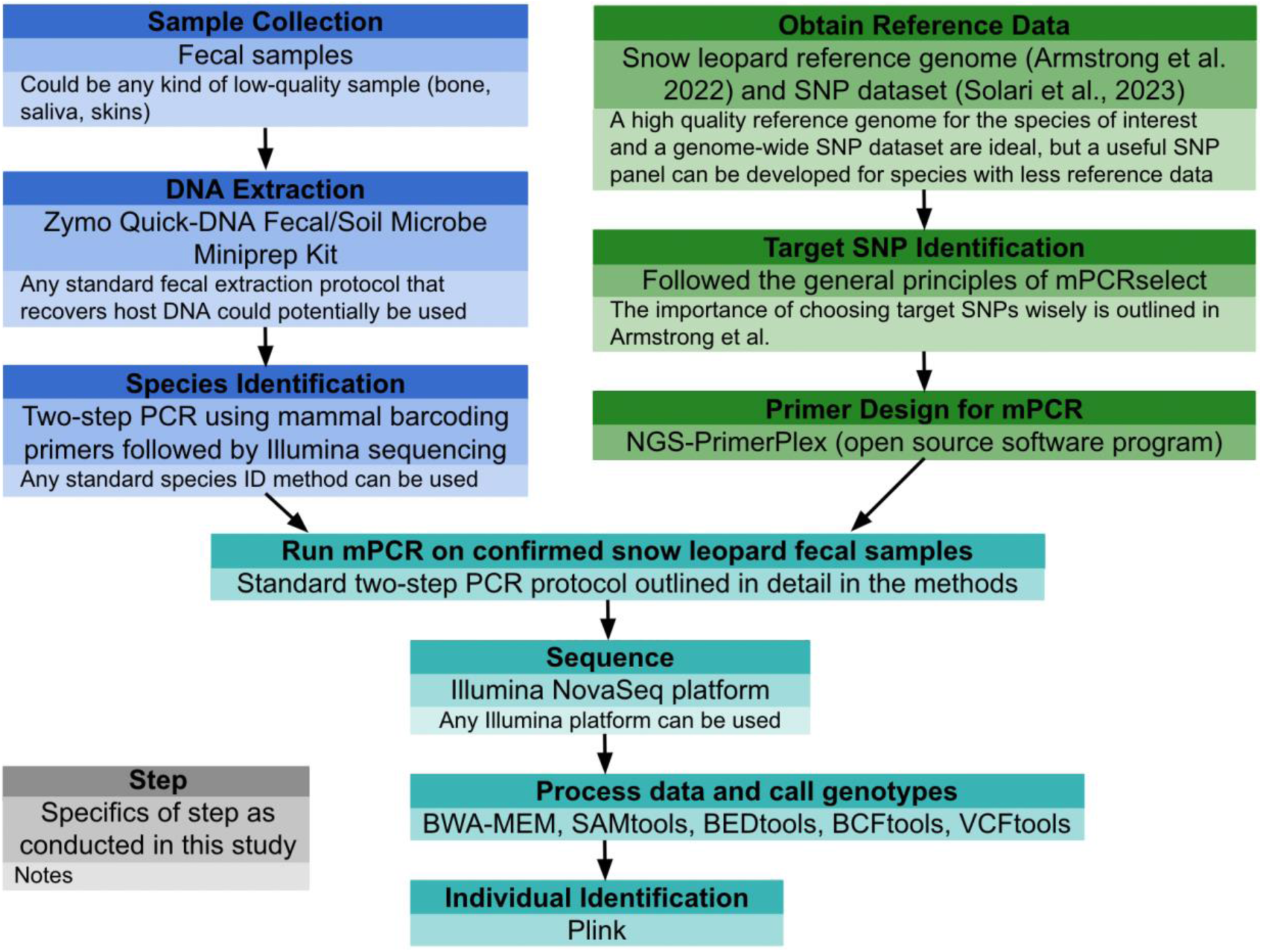
Schematic showing the steps taken to create a snow leopard SNP panel and conduct individual identification of fecal samples. The category of information provided in each portion of each box is indicated in the gray legend box.

### Species ID

We first assessed the species ID of the Pakistan fecal samples. Out of the 1,174 Pakistan fecal samples extracted and processed for species identification, 354 (30%) did not yield enough sequencing data for species identification (i.e., contained less than 50 reads using the mammal metabarcoding primers). Snow leopard was the second most common OTU. Fox was the most common OTU and dog/wolf (dogs and wolves are not distinguishable from the region amplified with MiMammal primers), lynx, and brown bear (*Ursus arctos*) were the third, fourth, and fifth most common OTU, respectively. OTU 14 was best matched to tiger (*Panthera tigris*). However, tigers are not present in this area and all samples with ten or more reads classified as OTU 14 also had reads classified as snow leopard. Thus, we reassigned OTU 14 to the closest matching species known to be in the area - snow leopard (97% identity).

Out of the 73 nulls (43 extraction nulls and 30 PCR nulls) that we ran, 18 yielded more than 50 reads. Of these 18, 12 were assigned to human, four to other mammals, and two to snow leopard.

We considered all samples with over 50 reads after all filtering and with over 51% of those reads assigned to snow leopard to be snow leopard scat and used them in downstream analyses. Using this cutoff, out of the 820 samples with sufficient sequencing data, we identified 235 fecal samples as snow leopard scat. Two extraction nulls passed this cut-off. Note that we ran all extraction nulls, regardless of species ID results, twice with the snow leopard SNP panel. All species ID read count results are provided in Table S1.

### SNP panel sequencing efficiency

In order to assess sequencing efficiency of the targeted loci, we determined the proportion of the total reads generated for each sample that mapped to target SNP regions. There was a wide spread in the amount of raw sequencing reads generated for each sample (Table S7) with a mean of ∼2.1 millions reads for each Pakistan run, ∼6.4 million for each SF Zoo run, ∼1.2 million for each null run, and ∼6.9 million for each non-snow leopard run. Of this data, a mean of 18% of reads (0–61%; SD = 21%) for each Pakistan run, 41% (0–61%; SD = 19%) of reads for each SF Zoo run, 15% (0.02–52%; SD = 20%) for each non-snow leopard run, and 0.002% (0–0.04%; SD = 0.004%) for each null run mapped to the target SNP regions. In all groups, the vast majority of spurious reads were low quality and did not map anywhere in the snow leopard genome (Table S7).

The mean depth of coverage for each target SNP per run after removing reads that mapped with a MQ of zero was 2,891X for the Pakistan runs, 21,917X for the SF Zoo runs, 0.084X for the nulls, 13,653X for other feline species, and 176X for canine species. Some target SNPs did sequence better than others; however, every target SNP sequenced successfully in at least some runs (Fig. S1).

### Genotyping success across samples

After calling SNPs and filtering for depth ≥6X, there was a mean of 65.4 successfully genotyped SNPs in each Pakistan run (n=609), 119.8 SNPs genotyped in each SF Zoo run (n=30), 0.06 SNPs genotypes in each extraction null (n=98), 0.6 SNPs genotypes in each PCR null (n=23), 95.4 SNPs genotyped in each non-snow leopard feline sample (n=12), and 4.6 SNPs genotyped in each canine samples (n=8) (Fig. 2A,B, Table S8).

**Fig. 2.**
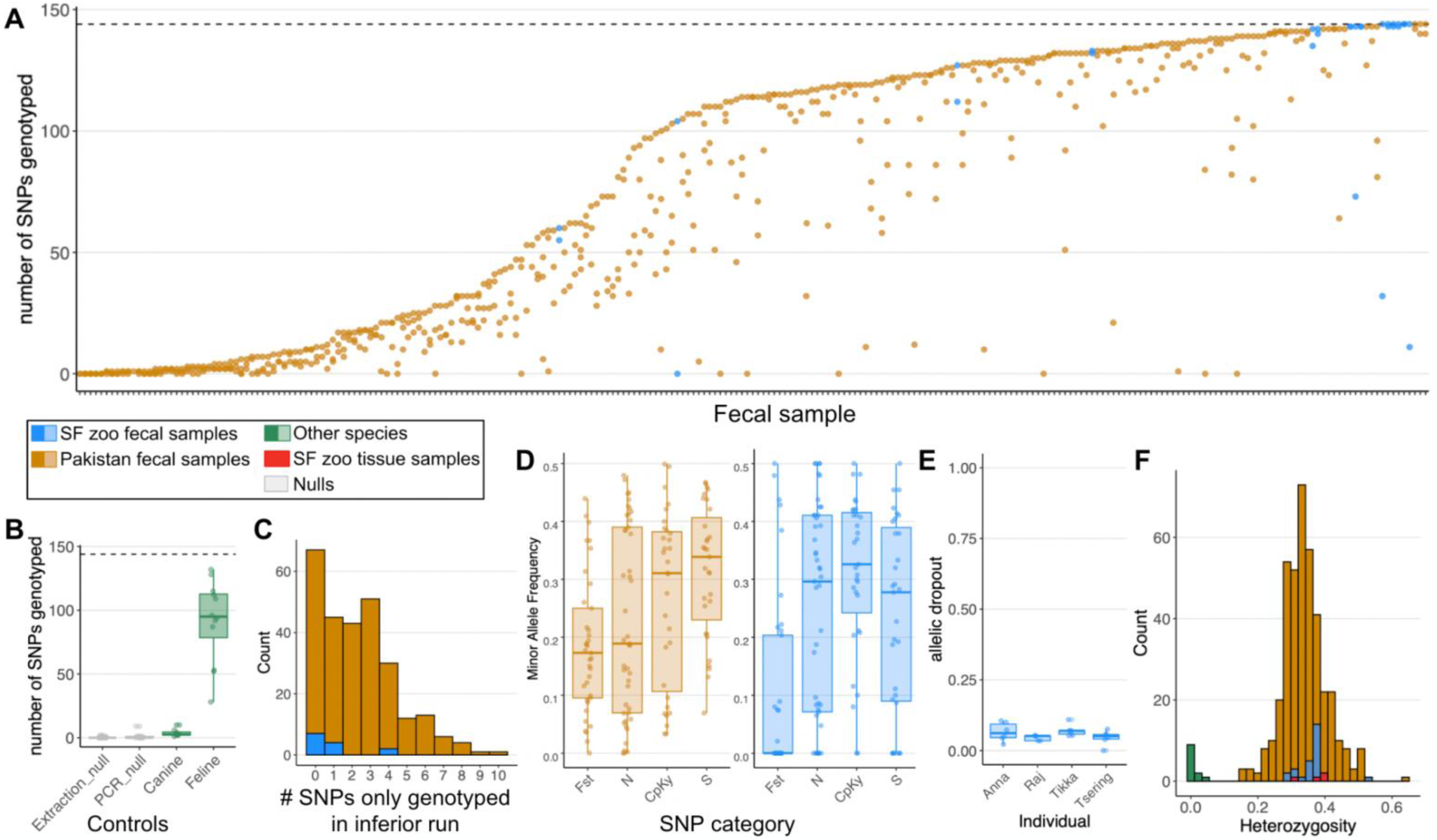
Snow leopard mPCR SNP panel performance summary. A) Scatter plot showing the number of SNPs genotyped in each replicate run of a sample to highlight the variability in run success among replicates. Samples are ordered by the number of SNPs successfully genotyped in their best run. SF Zoo fecal samples are shown in blue and Pakistan fecal samples are shown in dark orange. B) Box plots showing the number of SNPs genotyped in each control. Nulls are shown in gray and non-snow leopard fecal samples are shown in green. C) Histogram of the number of SNPs only genotyped in the inferior run when looking at all pairwise comparisons between replicate runs of the same sample showing a general lack in variability of which SNPs are successfully genotyped in replicate runs. D) The minor allele frequency for each SNP among all of the Pakistan samples (dark orange) and SF Zoo samples (blue) with SNPs separated by category (*F*_ST_ - SNPs chosen to target diversity between the groups, N - SNPs chosen to target diversity within the Northern group, CpKy - SNPs chosen to target diversity within the Captive/Kyrgyzstan group, and S - SNPs chosen to target diversity within the Southern group). E) Allelic dropout grouped by individual. Note that only four of the five SF Zoo individuals are shown here as only these four individuals had high-coverage WGS data available to serve as a ‘truth’ dataset. E) Histogram of heterozygosity for each sample, colored differently by group, showing that non-snow leopard samples are easily identifiable based on a lack of heterozygosity in target SNPs.

There were samples both in the SF Zoo sample set and the Pakistan sample set that showed high variability in genotyping success between replicates of the same run (Fig. 2A). Numerous samples from both the SF Zoo and Pakistan sample set were genotyped at all 144 SNPs; however, numerous samples from the Pakistan sample set did not sequence well in any replicate (Fig. 2A).

Out of the 121 nulls, 111 didn’t successfully genotype at any SNP, eight genotyped at one SNP, one genotyped at two SNPs, and one null genotyped at nine SNPs (Fig. 2B). Out of the eight canine fecal samples, all of them genotyped at ten SNPs or fewer; however, all of the 12 non-snow leopard feline samples successfully genotyped at 28 to 132 (mean 91.5) SNPs (Fig. 2B).

To assess the overlap between the SNPs genotyped in replicates of the same sample, we calculated the number of SNPs only genotyped in the inferior run for each pairwise comparison between replicate runs, where the inferior run is defined as the run with fewer successfully genotyped SNPs in each pairwise comparison. Among the 273 comparisons between replicate runs, we saw that the inferior run in the pair rarely added more than a few SNPs that were not already captured in the superior replicate (Fig. 2C).

### Information value of target SNPs

Solari et al. (2023) identified four genetically distinct snow leopard groups using WGS data from samples across the northwest of the species range. These four groups consist of samples from Russia and Mongolia (North group), samples from zoos and Kyrgyzstan (Captive/Kyrgyzstan group), samples from Pakistan, Afghanistan, and Tajikistan (South group), and a single sample from India which was genetically unique. Given this, we targeted four different categories of SNPs in this SNP panel – SNPs targeting diversity between geographic regions (*F*_ST_ SNPs), SNP targeting diversity within the Northern population (Northern SNPs), SNPs targeting diversity within the Captive/Kyrgyzstan population (Captive/Kyrgyzstan SNPs), and SNP targeting diversity within the Southern population (Southern SNPs). In order to assess how informative the target SNPs were in our sample sets, we calculated the minor allele frequency for each SNP in each sample set.

We know that the snow leopards that we sampled feces from in Pakistan are likely part of the Southern group and SF Zoo snow leopards are part of the Captive/Kyrgyzstan group. Thus, we expected SNPs targeting diversity within the Southern group to be most variable, and thus informative, among the Pakistan samples and the SNPs targeting diversity within the Captive/Kyrgyzstan group to be the most variable among the SF Zoo samples. This is indeed what we see in Fig. 2D. However, we also found SNPs in the other categories to be highly variable within our sample sets, showing that SNPs not specifically targeted for individual ID within a population can be highly valuable, especially in species like snow leopard with low *F*_ST_ values between groups (Solari et al. 2023).

We observe many more non-variable SNPs in the SF Zoo sample set (40 non-variable SNPs, only two of which were from the Captive/Kyrgyzstan category) compared to the Pakistan sample set (3 non-variable SNPs) (Fig. 2D). Given that the SF Zoo sample set consists of only five individuals and most of them are related, it is not surprising that we see less genetic variation among these samples than the larger Pakistan sample set.

### Genotyping accuracy and allelic dropout

We calculated genotyping accuracy and allelic dropout rate using SF Zoo fecal samples from individuals for which we had high-coverage WGS data (>40X) from paired tissue samples (n=24 runs representing 12 fecal samples from four individuals). We only retained SNP panel runs that were successfully genotyped at 20 SNPs or more. Twenty-three of the 24 runs passed this threshold. Across these 23 runs, the mean genotyping accuracy at the SNP level was 97.6% (minimum = 93.3%, maximum = 100%), the mean genotyping accuracy at the allele level was 98.8% (minimum = 96.7%, maximum = 100%), and the mean allelic dropout rate was 0.057 (minimum = 0, maximum = 0.11) (Fig. 2E). Note that accuracy is likely to be lower in more degraded field-collected samples.

### Heterozygosity

We calculated the heterozygosity of each run that successfully genotyped at 20 or more SNPs. All of the snow leopard fecal samples from Pakistan and SF Zoo as well as the SF Zoo tissue samples had a heterozygosity between 0.15 and 0.64 and all non-snow leopard feline samples had a heterozygosity lower than 0.04 (Fig. 2F). This shows that SNP panel results generated from non-snow leopard samples are easily distinguishable from snow leopard samples based on a lack of heterozygosity.

### Identifying individuals from mPCR output

We first filtered out runs that yielded genotype calls for less than 20 of the target SNPs. This filtering step removed all nulls (n=121), all canine samples (n=8), two SF Zoo runs, and 211 Pakistan sample runs. Thus, we were left with 398 Pakistan sample runs representing 180 of the 235 snow leopard fecal samples (77%), 28 of the 30 SF Zoo runs representing all 15 SF Zoo fecal samples (100%), and all of the original 12 runs of the non-snow leopard feline species.

We made pairwise comparisons between each of the 28 SF Zoo runs remaining post filtering. The distribution of the proportion of alleles IBS among all of the pairwise comparisons showed a clear separation between self-self comparisons and comparisons between samples from different individuals. All pairwise comparisons between samples from the same individual had more than 95% of alleles IBS and all comparisons between samples from different individuals had less than 86% of alleles IBS (Fig. 3A). Comparisons between SF Zoo samples had a mean of 112 SNPs for comparison and all comparisons had at least 29 SNPs.

**Fig. 3.**
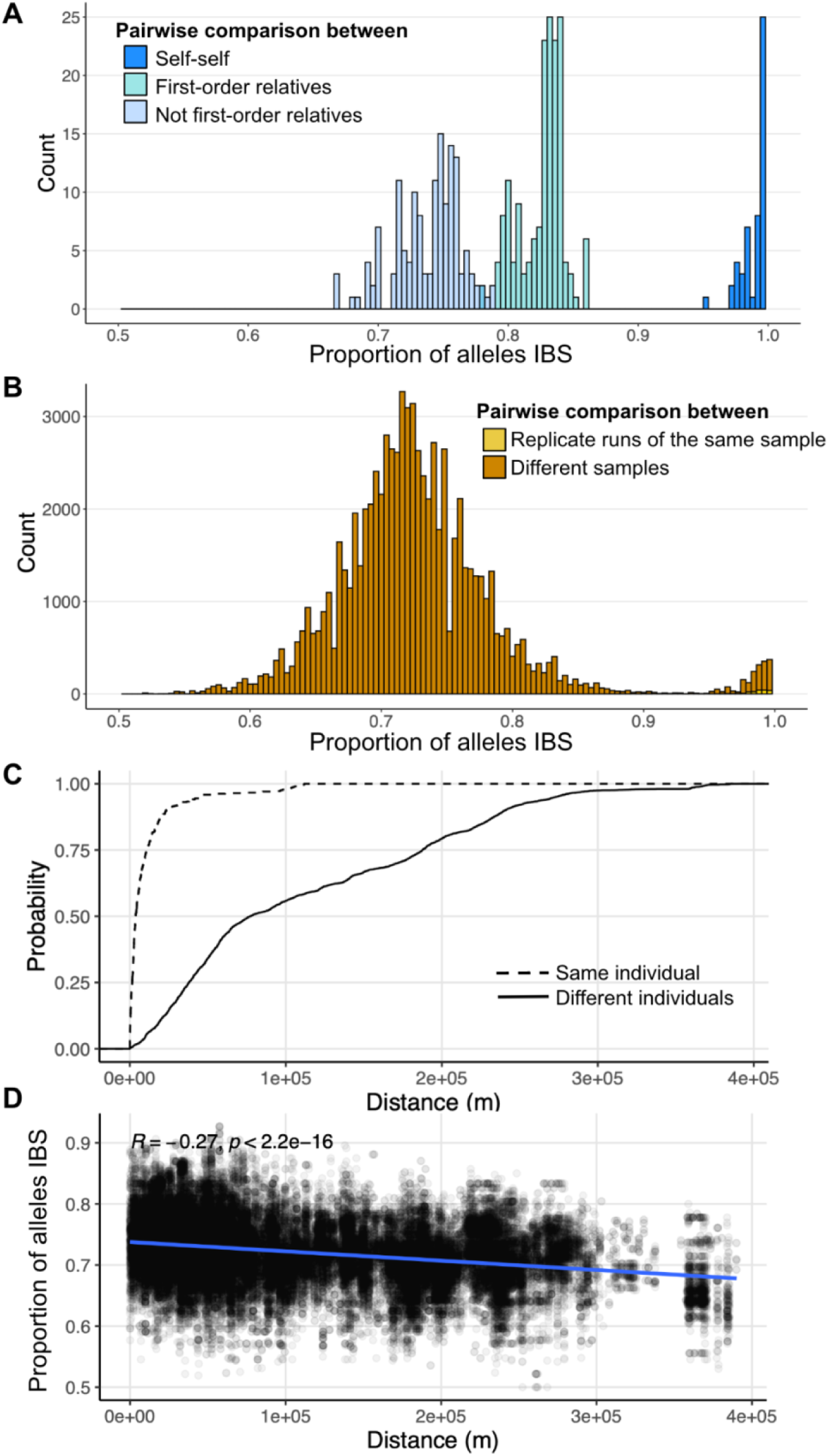
Assessments of individual ID and relatedness. A) Histogram showing the distribution of proportion of alleles IBS among all pairwise comparisons of runs within the SF Zoo sample set. All true relationships are known, so comparisons between self-self, between first-order relatives, and between non-first-order relatives are indicated with different shades of blue. B) Histogram showing the distribution of proportion of alleles IBS among all pairwise comparisons of runs within the Pakistan sample set. True relationships are unknown, so comparisons between replicate runs of the same sample are shown in yellow and comparisons between different fecal samples are shown in dark orange. C) Empirical Cumulative Distribution Function (ECDF) for distance between Pakistan samples from the same individual (proportion of alleles IBS > 0.95) and different individuals. D) Comparison of proportion of alleles IBS and distance among pairwise comparisons of Pakistan samples from different individuals. Each point is a pairwise comparison between runs from different individuals. The Pearson’s correlation coefficient and associated p-value are shown. In all plots, only pairwise comparisons consisting of 20 or more SNPs are included.

When plotting the distribution of the proportion of alleles IBS among all the pairwise comparisons between Pakistan sample runs made up of 20 SNPs or more (398 runs from 180 fecal samples), we found a bimodal distribution separating self-self comparison from self-non-self comparisons (Fig. 3B). We considered all comparisons with 95% or more alleles IBS to be from the same individual. Out of the 266 pairwise comparisons between replicates of the same sample that had 20 or more SNPs for comparison, eight did not meet this 95% cut off, all of which were comparisons with 35 or fewer SNPs.

Among all of the Pakistan runs, there were a total of 78,286 pairwise comparisons consisting of 20 SNPs or more of which 2,491 pairs had 95% of alleles or more IBS. This resulted in the identification of 35 unique snow leopards represented by 2 - 15 fecal samples and 21 unique snow leopards represented by one fecal sample, with no contradictory relationships (Fig. 4, Fig. S4, Table S6). ECDF analyses confirmed that two samples from the same individual were much more likely to be geographically close together than two samples from different individuals (Fig. 3D, Fig. S3).

**Fig. 4.**
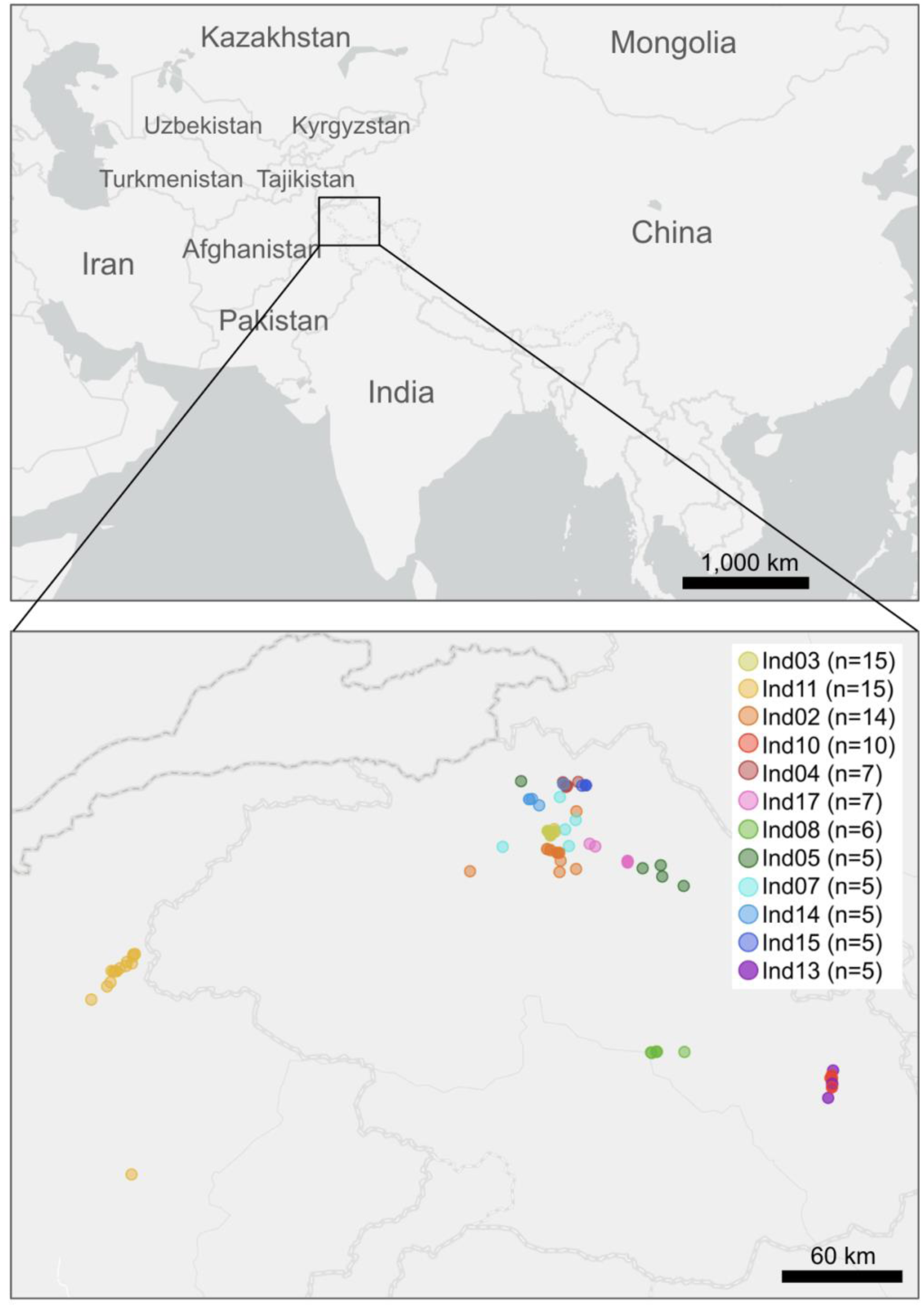
Map indicating Pakistan sampling area and the location of fecal samples from individuals represented by five or more fecal samples. The number of fecal samples from each individual is indicated. A similar map showing the location of fecal samples from individuals represented by four or fewer fecal samples is provided in the Supplementary (Fig. S4). Maps were created using ArcGIS software by ESRI and an ESRI basemap (ESRI, 2017).

### Assessing relatedness among samples

Out of the five SF Zoo individuals, four of the samples represented one family unit – mother, father, and two cubs. These known first-order relationships were largely identifiable using the proportion of alleles IBS with only minimal overlap between the two relatedness categories (Fig. 3A; note that it is not possible to distinguish full siblings from parent-offspring using IBS alone). All first-order relative pairs had between 78-86% of alleles IBS while unrelated pairs had less than 79% of alleles IBS. We found that first-order relationships did not overlap with non-first-order relationships when we increased the number of minimum SNPs used for pairwise comparison to 35 SNPs and were separated by a large gap in IBS values when increased to 75 SNPs (Fig. S2).

Among the Pakistan samples, for which there was no known relatedness data, we found a significant correlation between pairwise geographic distance between sample pairs and the proportion of alleles IBS (R = -0.27, p = 2.2^-16^, Fig. 3D) and an even more pronounced correlation when only looking at pairwise comparisons consisting of 75 SNPs or more (R = - 0.34, p = 2.2^-16^, Fig. S3). This indicates that our assumption that snow leopards are generally more related to individuals close by than those far away is correct.

### General geographic assignment

We included SNP panel results from one fecal sample from each SF Zoo snow leopard individual (n=5) and each unique Pakistan individual sampled (n=56) and conducted a PCA analysis with these samples and WGS data from across the range from Solari et al. (2023). The PCA analysis showed the fecal samples to group with the appropriate WGS samples – the SF Zoo samples group with the captive/Kyrgyzstan WGS data and the Pakistan samples group with the southern WGS samples (Fig. 5D).

**Fig. 5.**
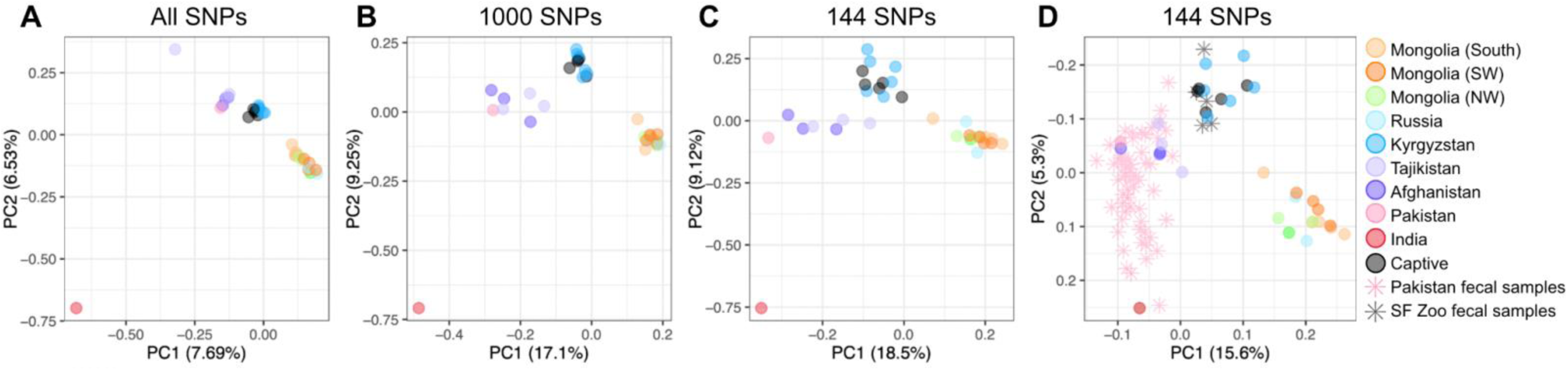
Principal component analysis (PCAs) showing genetic similarities between samples using different subsets of SNPs. A) PCA using all SNP data for all non-related individuals included in Solari et al. (2023) (n=33). B) PCA using only the 1,000 target SNPs for all the samples shown in panel A. C) PCA using only the 144 SNPs included in the SNP panel for the samples shown in panel A and B. D) PCA using only the 144 SNPs included in the SNP panel for the samples shown in previous panels as well as each unique fecal sample.

## Discussion

Here, we present an affordable protocol for SNP panel development and implementation that uses open source software for SNP panel design through data analysis, a streamlined lab protocol, and cost-effective Illumina sequencing technology (Fig. 1). We use this protocol to develop a SNP panel for individual ID in snow leopards from low-quality fecal samples which we show yields extremely accurate results, is robust to contamination (Fig. 2B), distinguishes non-target species (Fig. 2F), and offers insights into first order relatedness (Fig. 3A,D) and general geographic assignment (Fig. 5).

Across our two sample sets – fresh fecal samples from the SF Zoo and field-collected fecal samples of unknown age from Pakistan – 100% of fresh samples and 77% of field-collected samples yielded sufficient data for individual ID while none of the 121 null runs yielded enough data to make it past the initial filtering steps. For SF Zoo samples for which we had high-coverage WGS ‘truth’ data, a minimum of 96.7% of alleles were correctly called for all runs.

### SNP panel cost

There are numerous one-time investments that need to be made for a species in order to allow for SNP panel development – namely reference data consisting of a reference genome and WGS data for multiple individuals (Fig. 1). As indicated in Fig. 1, useful SNP panels can be made with varying degrees of reference data depending on the questions at hand. For example, if the goal of a project is individual ID within a small geographic area, then WGS from a few individuals from that area will likely suffice. However, if the goal is geographic assignment across the species range, then WGS from numerous individuals from each genetically distinct population will be necessary.

The primers used in running the SNP panel are another one-time investment that will likely generate enough primers to be shared across numerous labs and cost around $10 per primer pair from IDT. This includes SNP panel primers (PCR 1 primers) as well as indexing primers (PCR 2 primers). The other main wet-lab expense is the Master Mix from the Qiagen Multiplex PCR Kit (catalog # 206143). In the protocol described here, we used 14uL of this Master Mix per SNP panel run (10uL in PCR 1 and 4uL in PCR2) which comes out to around $2.20 per run just for the Master Mix.

Since this method takes full advantage of the most common and least expensive sequencing technology, sequencing costs can theoretically be trivial if one is able to purchase only a small portion of a sequencing lane. For example, at the sequencing facility used in this project, Admera Health (South Plainfield, New Jersey), one NovaSeq X Plus 25B lane generates 5-6.35 billion reads and costs $2,799. Assuming the lowest output for this lane (5 billion reads), one could generate 100X coverage for each of the 144 SNPs in this SNP panel for 125 runs for $1 (i.e. ∼1.8 million reads per $1). However, raw data processing results indicate that on average, only 19% of the raw data generated for low-quality fecal samples (Pakistan sample set) and 41% of the raw data generated for high-quality fecal samples (SF Zoo sample set) mapped well to the target regions. Additionally, samples from other species and nulls yielded a large number of raw reads (6.9 million and 1.2 million average reads, respectively) (Table S7). We also show that we greatly exceed the necessary level of sequencing here, so sequencing coverage, and thus the number of wasteful reads could easily be reduced. 100X coverage would be more than enough for our purposes and we exceeded that by over an order of magnitude in the Pakistan samples, and by more than two orders of magnitude for the SF Zoo samples. As seen in Table S7, at the cost of $1 for 1.8 millions reads, we spent $1-4 per run in this experiment. However, with additional effort put towards avoiding over-sequencing, we believe this cost could get much closer to the theoretical minimum of $1 per 125 runs.

### SNP panel recommendations

We show that some target SNPs did amplify better than others (Fig. S1); however, no target SNP failed in all runs (Fig. 2A). Also, we find that SNPs in categories focused on targeting diversity outside of the focal population are also highly variable (Fig. 2D), and thus informative to individual ID. This is because we expect highly variable SNPs in one snow leopard population to also be variable in other populations as the majority of SNPs in snow leopards are shared across populations (Solari et al., 2023). Thus, we recommend including all 144 SNPs in the SNP panel regardless of what focal population it is being run on.

We found that while some samples work consistently well or poorly across replicates, numerous samples had a great deal of variability in run success between replicates (Fig. 2A). Thus, we suggest that each sample be run at least twice even if there is no amplification success in the first run.

We also show that SNP panel results from non-snow leopards are easily identifiable such that there is no risk of mistakenly including data from the wrong species in the dataset if the panel is run on scat from a different species found in the snow leopard range. Canine samples (fox and dog/wolf) do not amplify well with the snow leopard SNP panel and were removed at the first filtering step due to insufficient amplification (Fig. 2B). Other feline samples (lynx, leopard, and leopard cat) did amplify well with the snow leopard SNP panel (Fig. 2B); however, we show that they are easily identified by the percent of heterozygous sites, as other feline species show almost no heterozygosity (0-2 heterozygous sites) in successfully genotyped SNPs (Fig. 2F). Thus, it is not pivotal to ensure that the SNP panel is not being run on non-snow leopard samples as these samples can be identified and removed post data generation.

We employed a cutoff of at least 95% of alleles IBS and 20 SNPs to classify two fecal samples as coming from the same individual. This cut off could be adjusted based on the specifics of the dataset, project goals, and how conservative researchers would like to be. Increasing the 20 SNP cutoff will likely result in throwing more samples out of the dataset and increasing or decreasing the 95% cutoff could result in more false negative matches or false positive matches, respectively.

### Relatedness among samples

Data for the SF Zoo sample set, for which first-order relatives are known, shows that first-degree relatives can be identified using this SNP panel (Fig. 3A) especially among sample comparisons with 75 or more SNPs (Fig. S2). These results suggest that among the Pakistan samples, sample pairs with a proportion of allele IBS of 0.80-0.94, especially those comparisons consisting of 75 or more SNPs (Fig. S3), are likely to be first-order relatives. Unknown characteristics of the Pakistan populations, such as levels of inbreeding, limit our ability to define an exact IBS range for first order relatives without reference data for that population; however, our panel offers hypotheses for relatedness.

### Conservation implications

The SNP panel that we present here, which enables individual ID of snow leopards from non-invasive fecal samples, can be used in numerous conservation contexts. Similarly to fecal-based individual ID using microsatellite markers, this data can be used to estimate local populations sizes and individual range sizes. Since the SNP panel is more cost and time effective than microsatellites, we hope that it could enable such assessments to be conducted repeatedly to allow conservation partners to track changes in snow leopard population size through time. Such temporal elements could also facilitate the assessment of snow leopard longevity in the wild. For example, among the samples included in this study, two individuals are represented by samples collected more than 4.5 years apart, offering some insight into longevity that could be extended with continued sampling efforts.

In addition to individual ID, we show that the SNP panel presented here provides information on general geographic origin (Fig. 5). Though only 37 population assignment SNPs were selected for this panel (*F*_ST_ SNPs), our results highlight how effective a SNP panel targeted solely geographic assignment of samples could be at offering precise origin assignments. This type of panel could be run on materials confiscated in the illegal wildlife trade (such as snow leopard skins) to identify where the animal originated such that mitigation efforts could be directed accordingly.

Lastly, we find that the SNP panel presented here can successfully distinguish first-degree relatives (Fig. 3A). If such insights into relatedness could be further validated with additional samples from known relatives and potentially strengthened with the inclusion of more SNPs for this explicit purpose, this could be extremely valuable to the conservation community. Accurate relatedness estimation could allow for assessments of reproductive rate, dispersal, etc.

More broadly, we present a protocol for SNP panel development and execution that is highly repeatable and uses only open-source software. This method offers a straightforward way to develop highly informative SNP panels for numerous use-cases for any species of interest for which the appropriate genomic reference data is available.

## Data Availability

SNP calling pipelines and analysis scripts are available on github (https://github.com/ksolari/SL_mPCR). The raw sequencing data are also available on github in vcf format.

## Code Availability

The code used for analyses in this project is available on the project’s github: https://github.com/ksolari/SL_mPCR

## Acknowledgements

This work was supported by funding from the Snow Leopard Trust and the Sabin Snow Leopards Grants Program. We would like to thank Gustaf Samelius and Koustubh Sharma from the Snow Leopard Trust for their help in securing funding for this work. We would like to thank the San Francisco Zoo, especially Adrian Mutlow and Stephanie Hees, for supplying the SF Zoo fecal samples used in this study. We extend our sincere gratitude to the provincial wildlife departments of Gilgit-Baltistan, Khyber-Pakhtunkhwa, and Azad Jammu and Kashmir for granting access to snow leopard habitats and supporting the sample collection process. We are also deeply appreciative of the local communities for their warm hospitality and invaluable assistance during sample collection. The dedicated team at the Snow Leopard Foundation played an instrumental role in the planning and execution of sample collection over several years. We would like to especially thank Nazakat Din, Muhammad Younas, Siraj Khan, and Raja Samandar Khan for their significant contributions.

