## Supplementary Table 4, 7 and 8 for "Next-generation snow leopard population assessment tool: multiplex-PCR SNP panel for individual identification from feces"

**Table S4. SF Zoo whole genome sequencing data stats**

| Individual | Depth of coverage | number of heterozygous sites from tissue WGS | Number of SNPs not genotyped from tissue WGS | percent heterozygous sites | Stud book number |
|---|---|---|---|---|---|
| Anna | 41.8 | 45 | 0 | 0.3125 | 117027 |
| Raj | 42.3 | 58 | 0 | 0.4028 | 117026 |
| Tikka | 48.1 | 57 | 0 | 0.3958 | 118008 |
| Tsering | 42.5 | 53 | 0 | 0.3681 | 118009 |
| Niya | 10.8 | 52 | 2 | 0.3662 | 111037 |

**Table S7. Read count summary statistics per run for each sample set**

| | Pakistan (n=609) | | SF Zoo (n=30) | | Other species (n=20) | | Null (n=121) | |
|---|---|---|---|---|---|---|---|---|
| | mean | SD | mean | SD | mean | SD | mean | SD |
| # raw reads | 2,082,201 | 2,683,831 | 6,437,453 | 4,546,193 | 6,914,276 | 2,451,279 | 1,196,095 | 2,150,961 |
| # reads with MQ>0 | 492,405 | 1,049,805 | 3,406,159 | 2,922,309 | 1,452,128 | 1,977,969 | 89,364 | 265,832 |
| # reads with MQ>0 mapping to amplicon | 410,223 | 969,837 | 3,168,710 | 2,740,913 | 1,196,752 | 1,921,925 | 27 | 137 |

SD = standard deviation, MQ = mapping quality

**Table S8. Number of SNPs (out of 144) successfully genotyped in each sample group**

| Sample group | mean | SD |
|---|---|---|
| Pakistan (n=609) | 63.6 | 53.4 |
| SF Zoo (n=30) | 118.3 | 43.4 |
| dog (n=4) | 4.3 | 4 |
| fox (n=4) | 3.3 | 1.9 |
| leopard (n=4) | 85 | 52.9 |
| leopard cat (n=4) | 102.2 | 9.7 |
| lynx (n=4) | 87.3 | 25.8 |
| Extraction nulls (n=98) | 0.06 | 0.3 |
| PCR nulls (n=23) | 0.56 | 1.88 |

SD = standard deviation
