## Supplementary Methods for "Next-generation snow leopard population assessment tool: multiplex-PCR SNP panel for individual identification from feces"

**Additional notes on steps taken to prevent contamination during SNP panel amplification:**

We did not visualize PCR1 product on a gel as this adds an additional opportunity for contamination of the PCR1 product (through an additional opening of the PCR1 product tube) before it is added to the PCR2 reaction. As mentioned in the main text, we found gel-visualization of both PCR1 and PCR2 products to be largely uninformative. The pre-PCR hood, where all PCR reactions were set up, had its own set of pipettes that never left the hood and which were UV-treated along with the rest of the hood before each use. When working in this pre-PCR hood, a lab coat was always worn to prevent any contaminants from clothing from getting into the hood. The lab coat worn while working in the hood was never exposed to the post-PCR space. As much as possible, the pre-PCR hood was not used after being in the post-PCR space - meaning that after going into the post-PCR space, we would wait to work in the pre-PCR hood until the following day, ideally wearing clothes that have not recently been in the post-PCR space.

When adding the PCR1 product to the PCR2 reaction in the post-PCR area, the following precautions were taken: the bench and pipette were bleached with 10% bleach, a new box of tips was used and the box was kept covered whenever not in use, the outside of the PCR1 product tubes were sprayed with 10% bleach, a paper towel was used to line the waste receptacle to try to cut down on aerosols, the waste container was placed at the opposite side of the bench from where the samples were being opened to prevent aerosols from getting close to the samples, each PCR1 tube was opened as briefly as possible, and each PCR2 tube was opened to add the PCR1 product as briefly as possible (each PCR2 tube was opened immediately before adding the PCR1 product and close immediately after).

Note that striptubes with individual attached lids were used for all species ID and SNP panel PCR1 and PCR2 reactions in order to allow for extra contamination-avoidance steps.
