## Supplementary File 1 for "Next-generation snow leopard population assessment tool: multiplex-PCR SNP panel for individual identification from feces"

Snow leopard SNP panel wet lab protocol

Katherine Solari (Petrov Lab, Stanford University)

v. 1.1 (2024-01-18)

#### **ATTENTION** - please contact Katie Solari if you are interested in running this protocol. I will send you the necessary primers and will answer any questions you may have.

#### NOTE - This protocol assumes that DNA has already been extracted from fecal samples and samples have already been confirmed as snow leopard. A suggested protocol for DNA extraction and species ID is provided in the appendix.

#### **Consumables**

| **Product** | **Manufacturer** | **Catalog #** |
| --- | --- | --- |
| 2x MasterMix from Qiagen Multiplex PCR Plus Kit | Qiagen | 206152 |
| PCR grade water |  |  |
| Bovine Serum Albumin (BSA) | Thermo Fisher | B14 |
| Snow leopard SNP panel primers | IDT | Will be supplied by Stanford |
| Unique dual indexing primers | IDT | Can be supplied by Stanford |
| PCR strip tubes with individual attached lids |  |  |
| 10% bleach | any |  |
| 70% ethanol | any |  |
| Zymo clean and concentrator-25 | Zymo | R1017 |

#### **Labware**

- pre-PCR hood with built-in UV (if you do not have one, alternatives are likely possible - contact me)
- 200uL and 10uL pipettes (these pipettes should live in the pre-PCR hood and never be removed)
- 10uL multichannel pipette - not essential, but is very nice to have for the addition of indexing primers (if primers are in plate format) during PCR2 set up (This pipette should live in the pre-PCR hood and never be removed)
- Small vortexer
- Small centrifuge for PCR strip tubes
- thermocycler

#### **Preparation**

- UV hood before use
- Wipe down hood with 10% bleach followed by 70% ethanol before use
- Place paper towels in the waste container to limit aerosoles
- Thaw and thoroughly mix fecal DNA

#### **Notes**

- Be sure to also process extraction nulls
- Be sure to include a PCR1 and PCR2 null as indicated in the procedure

#### **Procedure (for n number of samples)**

1. In a pre-PCR hood make the PCR1 master mix consisting of:
   a. 10 x (n+1) uL 2x MasterMix from Qiagen Multiplex PCR Plus Kit
   b. 1 x (n+1) uL water
   c. 1 x (n+1) uL BSA
   d. 4 x (n+1) uL of Snow Leopard SNP panel primer pool at 10uM
2. Distribute 15 uL of this master mix into n+1 PCR tubes (ensure you are using PCR strip tubes with individual attached lids) and close all lids.
3. Add 4uL of extracted fecal DNA to each tube. Only have one DNA tube in the hood at a time and only open one PCR strip tube at a time. Add 4uL of water to the PCR 1 null (sample n+1).
4. Ensure all strip tube lids are closed and remove them from the hood.
5. Briefly vortex the samples and pulse spin down
6. Place the tube in a thermocycler that is ideally in a different room, lab, or building than your pre-PCR hood. Run the following program:
   a. 95°C for 5 min
   b. 95°C for 30 sec
   c. 60°C for 90 sec
   d. 72°C for 30 sec
   e. Repeat steps (b–d) 40 times
   f. 68°C for 10 min
   g. Hold at 4°C
7. Ideally on a different day, in the pre-PCR hood, prepare the PCR2 Master Mix consisting of:
   a. 4 x (n+2) uL 2x MasterMix from Qiagen Multiplex PCR Plus Kit
   b. 3 x (n+2) uL water
8. Distribute 7 uL of this master mix into n+2 PCR tubes and close all lids.
9. Add 1uL of each of the forward and reverse indexing primers at 10mM to each PCR tube. Be sure to add a unique combination of indexing primers to each well and note which indexes have been added.
10. Close tubes, remove them from the pre-PCR hood and bring them to your post-PCR space where you currently have the PCR1 product from step 6.
11. Prepare your post-PCR lab space:
    a. bleach the bench you are working on
    b. Bleach the pipette you will be using to transfer PCR1 product
    c. Thoroughly cover PCR1 product tubes in bleach before opening
12. Add 1uL of PCR1 product to each PCR2 master mix. Be sure to open both the PCR1 tube and the PCR2 tube as briefly as possible. Do not open or add anything to the PCR2 null (well # n+2)
13. Briefly vortex the samples and pulse spin down
14. Bleach the PCR machine and place the tubes in the thermocycler. Run the following program:
    a. 95°C for 5 min
    b. 95°C for 30 sec
    c. 62°C for 90 sec
    d. 72°C for 30 sec
    e. Repeat steps (b–d) 12 times
    f. 68°C for 10 min
    g. Hold at 4°C
15. Pool 2uL of PCR2 product from each sample and clean this pool using a Zymo clean and concentrator-25 kit following the manufacturer's protocol.
16. This is your final library. Sequence it on an Illumina MiSeq machine using a 1x250 bp configuration. Aim for more than 30,000 reads per sample.

#### **Additional notes on preventing contamination:**

- We recommend that you do not visualize PCR1 or PCR2 products on a gel. We have found gel visualizations to be uninformative regarding amplification success and thus this is simply an additional opportunity for contamination.
- Keep all fecal DNA extractions in your designated pre-PCR space and never open these tubes outside of a hood.
- Keep all PCR products in the post-PCR space and NEVER bring them to the pre-PCR space.
- Maintain a unidirectional flow of materials, such that nothing from the post-PCR lab space comes back to the pre-PCR lab space.
  - Keep all materials (consumables, reagents, gloves, papers) used in the post-PCR space in the post-PCR space. Avoid bringing anything from the post-PCR space to the pre-PCR space. NEVER bring anything from the post-PCR space into the pre-PCR hood.
- Treat the pre-PCR hood with UV light, bleach and ethanol before each use.
- Attempt to physically separate your pre-PCR space and your post-PCR space as much as possible (separate rooms, floors, or buildings)
- The pre-PCR hood, where all PCR reactions are set up, should have its own set of pipettes that never leave the hood and which are UV-treated along with the rest of the hood before each use.
- When working in the pre-PCR hood, a lab coat should always be worn to prevent any contaminants from clothing from getting into the hood. The lab coat worn while working in the hood should never be exposed to the post-PCR space
- To the extent possible, do not use the pre-PCR hood after being in the post-PCR space.
  - After going into the post-PCR space, if possible, wait to work in the pre-PCR hood until the following day, ideally wearing clothes that have not recently been in the post-PCR space.
- When adding the PCR1 product to the PCR2 reaction in the post-PCR area:
  - Bleach the bench and pipette with 10% bleach
  - Use a new unopened box of tips and keep the box closed/covered when not in use.
  - Spray the outside of the PCR1 product tubes with 10% bleach
  - Use a paper towel to line the waste receptacle to try to cut down on aerosols
  - Place the waste container at the opposite side of the bench from where the samples are being opened to prevent aerosols from getting close to the samples
  - Only open one PCR1 tube at a time and keep it open as briefly as possible
  - Only open one PCR2 tube at a time and keep it open as briefly as possible (open each PCR2 tube immediately before adding the PCR1 product and close immediately after)

### **Appendix**

### **Supplementary protocol 1 - DNA extraction**

#### *Note that numerous DNA extraction methods can be used. If your lab has already extracted DNA from fecal samples for microsatellite analyses, the same DNA extraction method should work well for this protocol as well.

#### **Consumables**

| **Product** | **Manufacturer** | **Catalog #** |
| --- | --- | --- |
| Zymo Quick-DNA Fecal/Soil Microbe Miniprep Kit (50 samples) | Zymo | D6010 |
| 1.5mL DNA LoBind tubes | Eppendorf | 022431021 |
| Razor blades | any |  |
| Sterile weighing boats (large enough to hold a fecal sample) | Any ([example](https://www.fishersci.com/shop/products/fisherbrand-polystyrene-antistatic-weighing-dishes-7/08732115)) |  |
| 10% bleach | any |  |
| 70% ethanol | any |  |

#### **Labware**

- 1000uL and 200uL pipettes
- Hood with built-in UV (if you do not have one, alternatives are likely possible - contact me)
- Beadbasher - Ideally one of the models recommended on page 8 of the Zymo Quick-DNA Fecal/Soil Microbe Miniprep handbook (<https://files.zymoresearch.com/protocols/_d6010_quick-dna_fecalsoil_microbe_miniprep_kit.pdf>)
- 2mL tube centrifuge that goes up to 10,000xg
- Small vortexer

#### **Preparation**

- UV hood before use
- Wipe down hood with 10% bleach followed by 70% ethanol before use
- Place paper towels in the waste container to limit aerosoles

#### **Notes**

- Be sure to include an extraction null with every set of samples.
- We recommend not processing more samples than will fit in the centrifuge you are working with.

#### **Procedure**

1. Only process one fecal sample at a time in the hood:
   a. Move the sample to a new sterile weighing boat. Use tweezers and a new sterile razor blade to scrape <150 mg (pea sized amount) of material from the outside of the feces. Put the fecal sample back in its original container and then transfer the scraped fecal material from the weigh boat to the 2mL beadbashing tube (included in the Zymo kit).
   b. Try to get material from the outside of the feces as this is where the snow leopard DNA is most concentrated.
   c. Wipe down the hood space with bleach and ethanol between samples.
2. Follow the manufacturer’s protocol for the Zymo DNA extraction kit.
   a. Note- we homogenized samples using the biospec mini beadbeater 96 for 5 minutes
3. Elute the final DNA in a 1.5mL LoBind tube for long-term storage.

#### **Supplementary protocol 2 - Species Identification**

#### *Note that numerous species identification methods can be used. The main goal here is to confirm that fecal samples are from snow leopards before running the SNP panel.

#### **Consumables**

| **Product** | **Manufacturer** | **Catalog #** |
| --- | --- | --- |
| AmpliTaq Gold 360 Master Mix | Thermo Fisher | 4398881 |
| PCR grade water | any |  |
| Bovine Serum Albumin (BSA) | Thermo Fisher | B14 |
| MiMammal forward primer with sequencing adapter - TCGTCGGCAGCGTCAGATGTGTATAAGAGACAGNNNNNNGGGTTGGTAAATTTCGTGCCAGC | IDT |  |
| MiMammal reverse primer with sequencing adapter - GTCTCGTGGGCTCGGAGATGTGTATAAGAGACAGNNNNNNCATAGTGGGGTATCTAATCCCAGTTTG | IDT |  |
| Unique dual indexing primers | IDT | Can be supplied by Stanford |
| 10% bleach | any |  |
| 70% ethanol | any |  |
| Zymo clean and concentrator-25 | Zymo | R1017 |

#### **Labware**

- pre-PCR hood with built-in UV (if you do not have one, alternatives are likely possible - contact me)
- 200uL and 10uL pipettes (These pipettes should live in the pre-PCR hood and never be removed)
- 10uL multichannel pipette - not essential, but is very nice to have for the addition of indexing primers (if primers are in plate format) during PCR2 set up (This pipette should live in the pre-PCR hood and never be removed)
- Small vortexer
- Small centrifuge for PCR strip tubes
- thermocycler

#### **Preparation**

- UV hood before use
- Wipe down hood with 10% bleach followed by 70% ethanol before use
- Place paper towels in the waste container to limit aerosoles

#### **Notes**

- Be sure to also process extraction nulls
- Be sure to include a PCR1 and PCR2 null

#### **Procedure (for n number of samples)**

1. In a pre-PCR hood make the PCR1 master mix consisting of:
   a. 5 x (n+1) uL AmpliTaq Gold 360 Master Mix
   b. 2 x (n+1) uL water
   c. 0.5 x (n+1) uL BSA
   d. 0.5 x (n+1) uL Forward Primer at 10mM
   e. 0.5 x (n+1) uL Reverse Primer at 10mM
2. Distribute 8 uL of this master mix into n+1 PCR tubes (ensure you are using PCR strip tubes with individual attached lids) and close all lids.
3. Add 2uL of extracted fecal DNA to each tube. Only have one DNA tube in the hood at a time and only open one PCR strip tube at a time. Add 2uL of water to the PCR 1 null (sample n+1).
4. Ensure all strip tube lids are closed and remove them from the hood.
5. Briefly vortex the samples and pulse spin down
6. Place the tube in a thermocycler that is ideally in a different room, lab, or building than your pre-PCR hood. Run the following program:
   a. 95°C for 10 min
   b. 95°C for 30 sec
   c. 60°C for 30 sec
   d. 72°C for 30 sec
   e. Repeat steps (b–d) 32 times
   f. 72°C for 7 min
   g. Hold at 4°C
7. Ideally on a different day, in the pre-PCR hood, prepare the PCR2 Master Mix consisting of:
   a. 5 x (n+2) uL AmpliTaq Gold 360 Master Mix
   b. 2.5 x (n+2) uL water
8. Distribute 7 uL of this master mix into n+2 PCR tubes and close all lids.
9. Add 0.5uL of each of the forward and reverse indexing primers at 10mM to each PCR tube. Be sure to add a unique combination of indexing primers to each well and note which indexes have been added.
10. Close tubes, remove them from the pre-PCR hood and bring them to your post-PCR space where you currently have the PCR1 product from step 5.
11. Prepare your post-PCR lab space:
    a. bleach the bench you are working on
    b. Bleach the pipette you will be using to transfer PCR1 product
    c. Thoroughly cover PCR1 product tubes in bleach before opening
12. Add 1.5uL of PCR1 product to each PCR2 master mix. Be sure to open both the PCR1 tube and the PCR2 tube as briefly as possible. Do not open or add anything to the PCR2 null (well # n+2)
13. Briefly vortex the samples and pulse spin down
14. Bleach the PCR machine and place the tubes in the thermocycleer. Run the following program:
    a. 95°C for 10 min
    b. 95°C for 30 sec
    c. 60°C for 30 sec
    d. 72°C for 10 sec
    e. Repeat steps (b–d) 12 times
    f. 72°C for 7 min
    g. Hold at 4°C
15. Run all extraction and PCR nulls and a subset of the PCR2 product on a 1% gel to monitor for contamination.
16. Once ensuring no contamination, pool 2uL of PCR2 product from each sample and clean this pool using a Zymo clean and concentrator-25 kit following the manufacturer's protocol.
17. This is your final library. Sequence it on an Illumina MiSeq machine using a 2x250 bp configuration. Aim for more than 250 reads per sample.
