## Supplementary Figures for "Next-generation snow leopard population assessment tool: multiplex-PCR SNP panel for individual identification from feces"

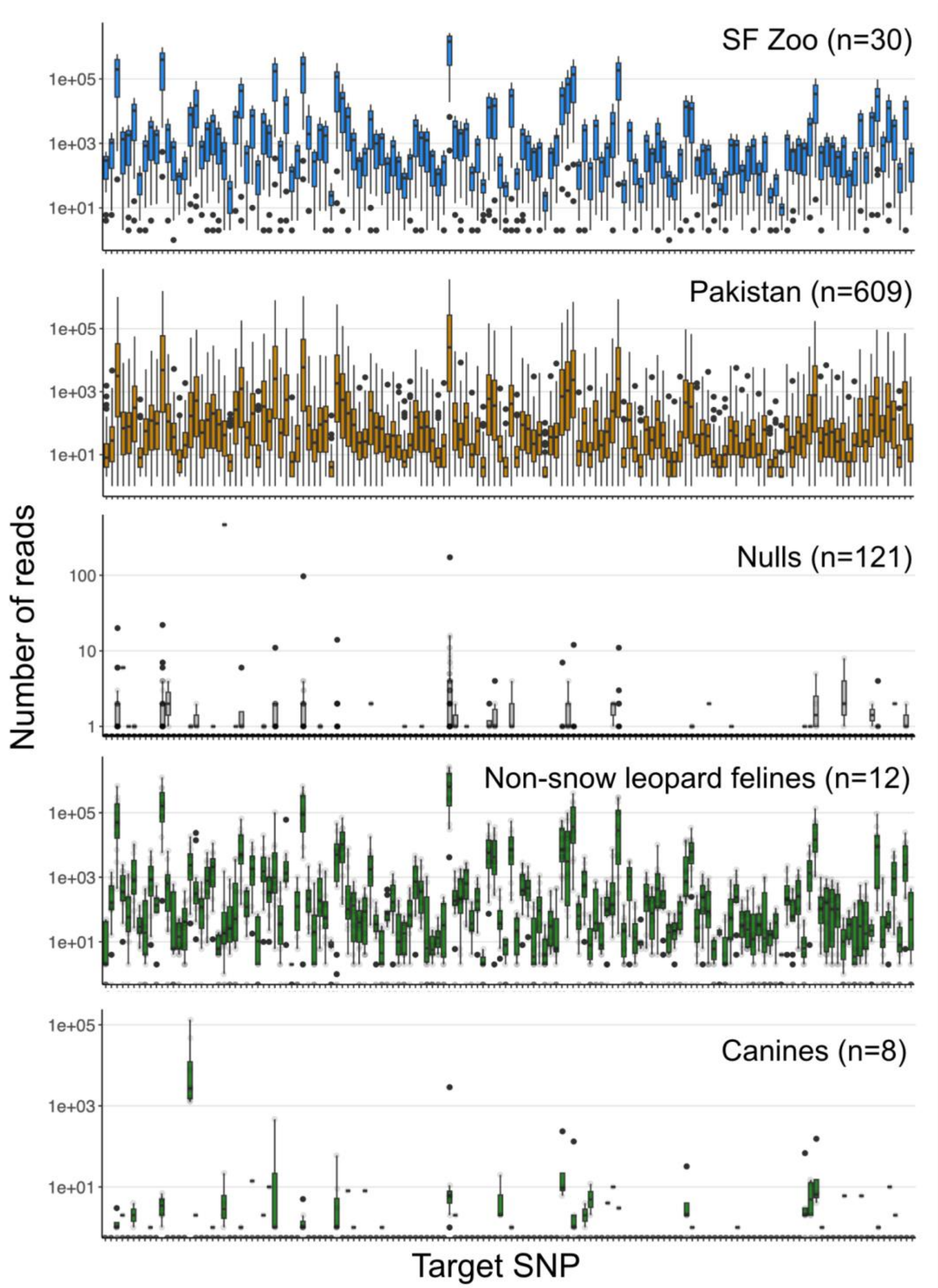


**Fig. S1. Box plots showing the number of reads per target SNP in each sample group.** The sample group and number of runs in each group is indicated above each plot. For nulls and non-snow leopard samples, raw data points are plotted faintly along with box plots. In all plots, the lower and upper edges of the boxes correspond to the first and third quartiles and the whiskers extend to the lowest/highest value that is no further than 1.5*IRQ (inter-quartile range) from the box. Points falling further than 1.5*IRQ from the box are plotted individually.

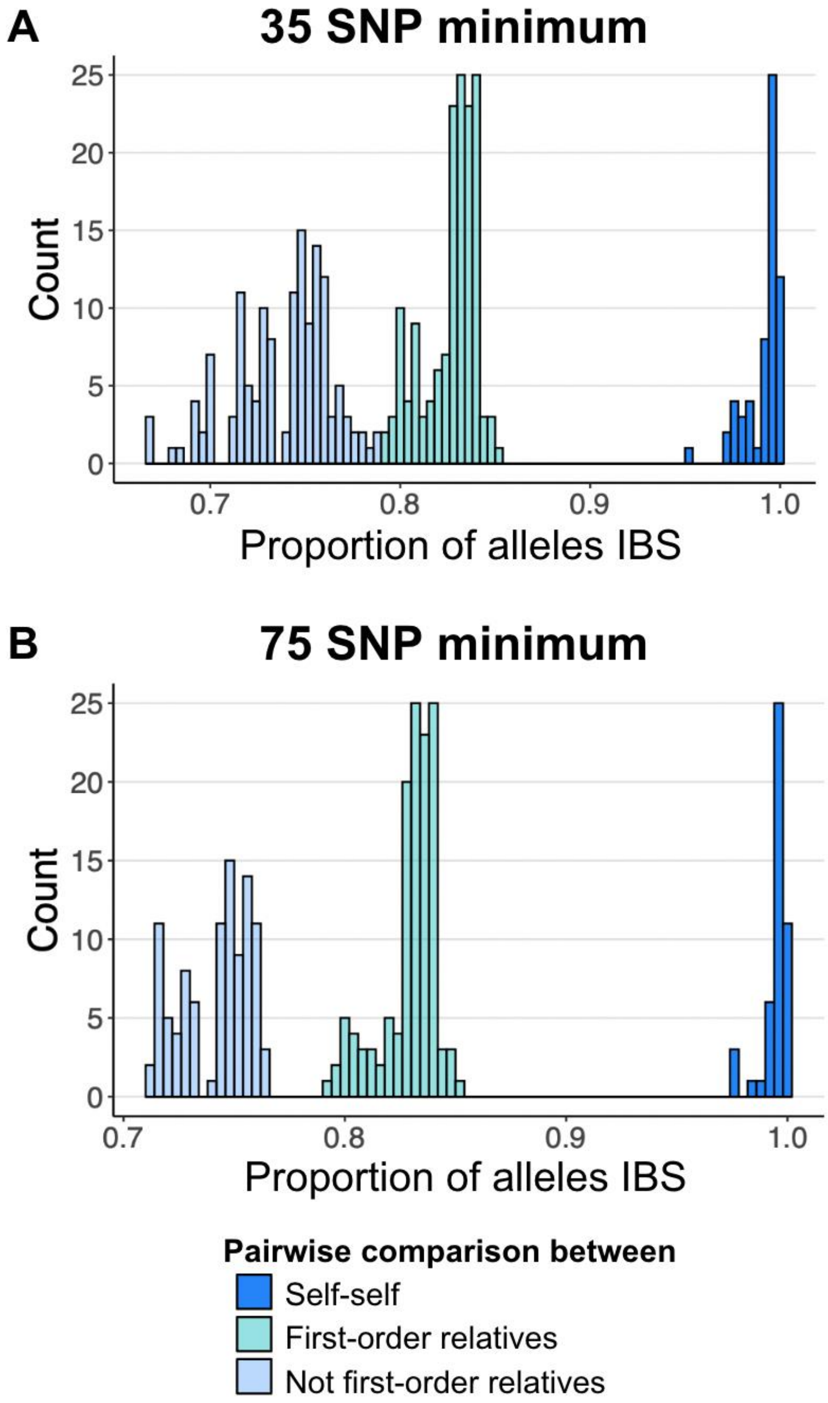


**Fig. S2. Histogram showing the distribution of proportion of alleles IBS among all pairwise comparisons within the SF Zoo sample set.** A) Histogram of pairwise comparisons consisting of 35 SNPs or more. B) Histogram of pairwise comparisons consisting of 75 SNPs or more. All true relationships are known, so comparisons between self-self, between first-order relatives, and between non-first-order relatives are indicated with different shades of blue.

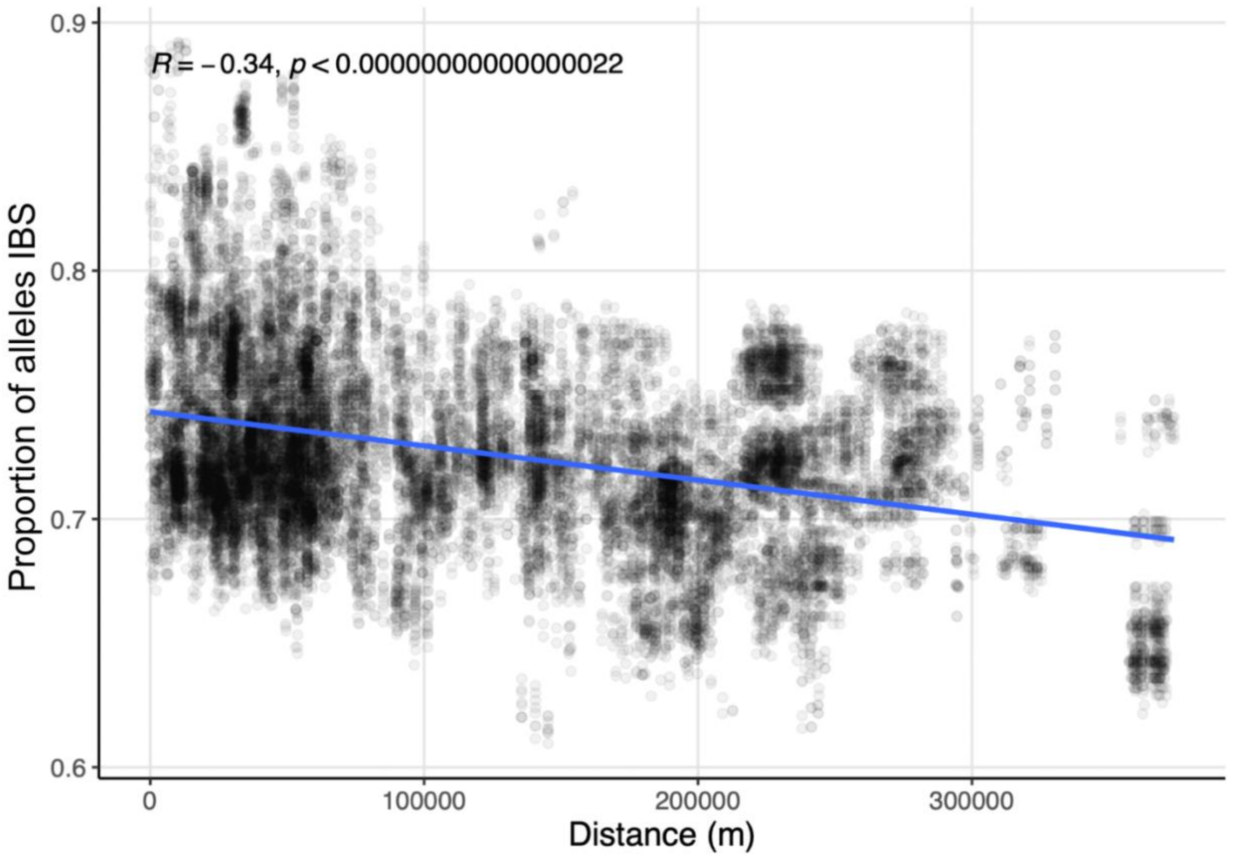


**Fig. S3.** Comparison of proportion of alleles IBS and distance among pairwise comparisons consisting of 75 SNPs or more. Each point is a pairwise comparison between two fecal samples from different individuals. The Pearson's correlation coefficient and associated p-value are shown.

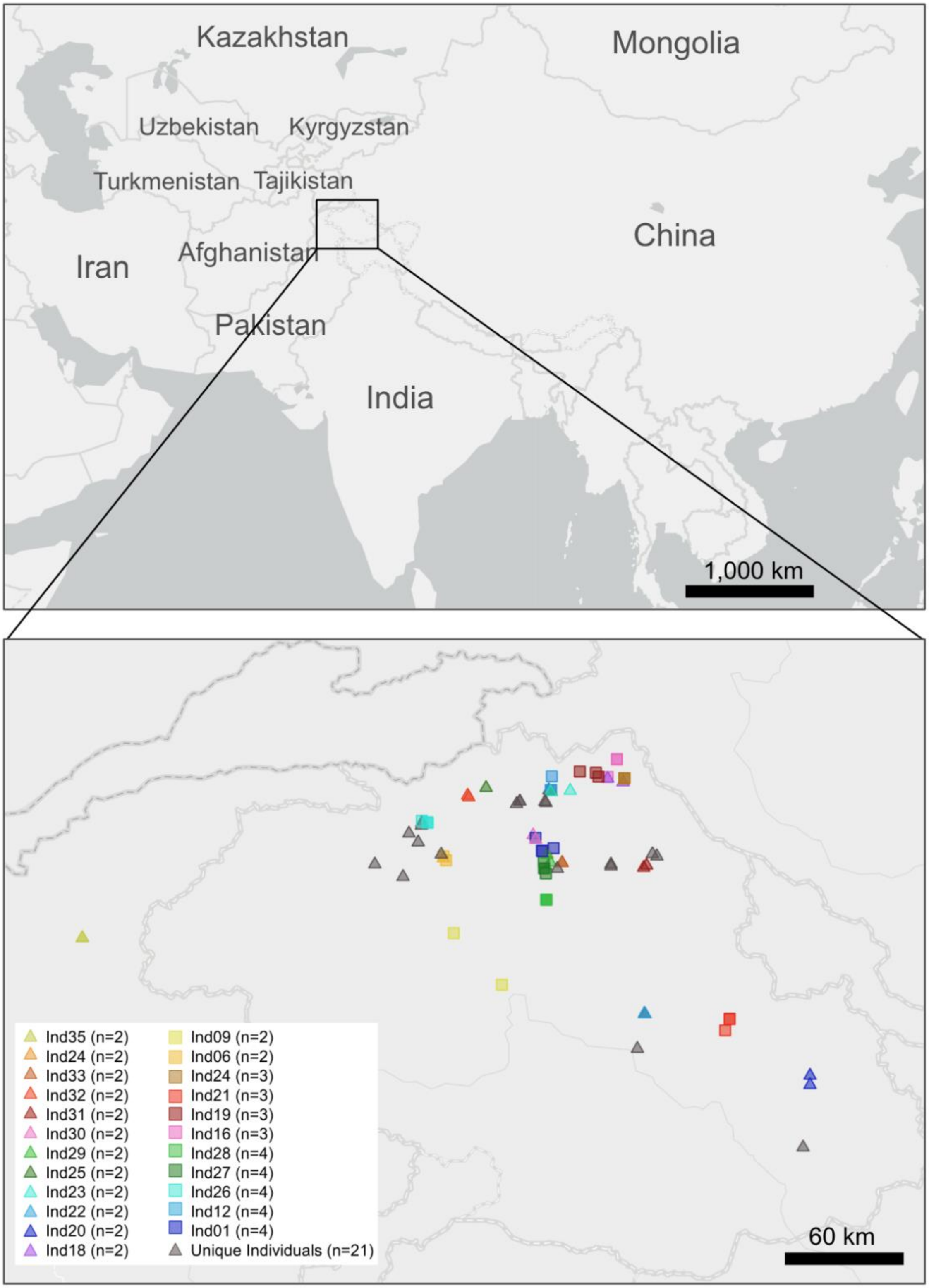


**Fig. S4. Map indicating Pakistan sampling area and the location of fecal samples from individuals represented by 4 or fewer fecal samples.** The number of fecal samples from each individual is indicated. Maps were created using ArcGIS software by ESRI and ESRI basemap(ESRI, 2017).
